# Toward Interpretable Digital Biomarkers of Walking and Reaching in Parkinson’s Disease

**DOI:** 10.1101/2022.04.23.489070

**Authors:** Jihye Ryu, Elizabeth Torres

## Abstract

Multimodal digital data registered with wearable biosensors has emerged as highly complementary of clinical pencil-and-paper criteria, offering new insights in ways to detect and diagnose various aspects of Parkinson’s disease. A pressing question is how to combine both the clinical knowledge of PD and the new technology to create interpretable digital biomarkers easily obtainable with off-the-shelf technology. Several challenges concerning disparity in biophysical units, anatomical differences across participants, sensor positioning, and sampling resolution are addressed in this work, along with identification of optimal parameters to automatically differentiate patients with PD from controls. We combine data from a multitude of biosensors registering signals from the central (EEG) and peripheral (magnetometry, kinematics) nervous systems, inclusive of the autonomic nervous system (EKG), as the participants perform natural tasks requiring different levels of intentional planning and automatic control. We find that magnetometer data during walking, across a variety of amplitude and timing signals provide optimal separation of PD from neurotypical controls. We conclude that considering multimodal signals, while differentiating across levels of intent in natural actions can be revealing of important features of PD that otherwise escape the naked eye. Further we add that clinical criteria combined with such optimal digital parameter spaces offer a far more complete picture of PD than using either one of these pieces of data alone.

## 1. Introduction

The advent of wearable biosensors with research grade sampling resolutions has opened new avenues of inquiry in basic clinical research, offering the possibility of unobtrusive data registration in remote telemedicine settings. While these new advances have the potential to expand the uses of such techniques at scale, beyond the confines of the laboratory, they also pose new challenges to data analysis and interpretation. Often, wearable biosensors output a multiplicity of data streams amenable to various interpretations, so one question is how to combine their information most efficiently to *e.g*., maximally differentiate features of a given disorder and highlight their departure from stages of natural aging. One possible way to leverage the richness of information in data from wearables is by developing experimental assays that evoke and combine different degrees of mental intent in action generation, with parameter identification revealing maximal differentiation across a cohort of participants. This can be achieved by discovering kinematic and biorhythm-based parameter spaces that maximally separate participants into strata revealing specific motor differences.

In the case of participants with Parkinson’s disease (PPD), this approach could serve to differentiate various aspects of motor performance from those of neurotypical participants, aging without neurological issues. For example, tasks such as pointing, and walking differ in their level of required guidance to motor control. Pointing is usually centrally guided, requiring precise (often visual) guidance to upper body limbs control and cortically based planning (*1, 2*). In contrast, walking involves more automated functions, centrally and peripherally guided by loops involving central pattern generators (CPGs) in the spinal cord, along with subcortical and cerebellar brain structures (*3*). Both sets of actions require the solution of Bernstein’s degrees of freedom (DOF) problem (*4*), which can be partly appreciated through kinematic and muscle synergies that recruit and release DOF in the corresponding intrinsic joint angle (*5, 6*) and muscle spaces (*7, 8*) that the brain must navigate throughout the motion. Likewise, both sets of actions have different levels of moment-to-moment variability (*9–11*) which must necessarily impact the types of sensory feedback returning to the brain, along with the latencies at which such feedback arrives to the spinal cord and brain centers (*3*).

Multimodal data streams may have different physical units which provide different possible interpretations of the parameters under consideration, in relation to their ranges and summary population statistics. They also provide information on the individual ranges, but only when the parameters have been properly standardized and brought to a unitless scale that permits comparison across often highly heterogeneous cohorts of participants.

In this work, we present various data types, data transformations that nevertheless preserve the original physical ranges, and parameter spaces derived from the biosensor’s multimodal data. These derivative data account for possible allometric effects inherently present in diverse cohorts of participants with differing anatomical features. The data allows us to consider parameter spaces inclusive of all participants in a random draw of the population and seek automatic stratification of the cohort into *e.g.,* neurotypicals and PDPs. Furthermore, we present methods of analyses that consider disparities in patterns of moment-to-moment variability, thus seeking a stochastic characterization that optimally separates such subgroups in various parameter spaces (**Figure 1** motivates our quest here.) This work considers multiple parameters and parameter spaces and identifies those which maximally separate members of the cohort, in a statistical and stochastic sense amenable for personalized approaches leading to clinically meaningful strata. We coin these parameters *interpretable digital biomarkers*. They are derived from continuous streams of multimodal digital data and self-emerge from the inherent variability unique to each person. Yet, they conserve the clinical interpretation that defines the disorders in the first place. While these methods aim at closing the gap between clinical and computational frameworks, they also reveal new information that may escape the naked eye of an expert during clinical evaluations.

**Figure 1.**
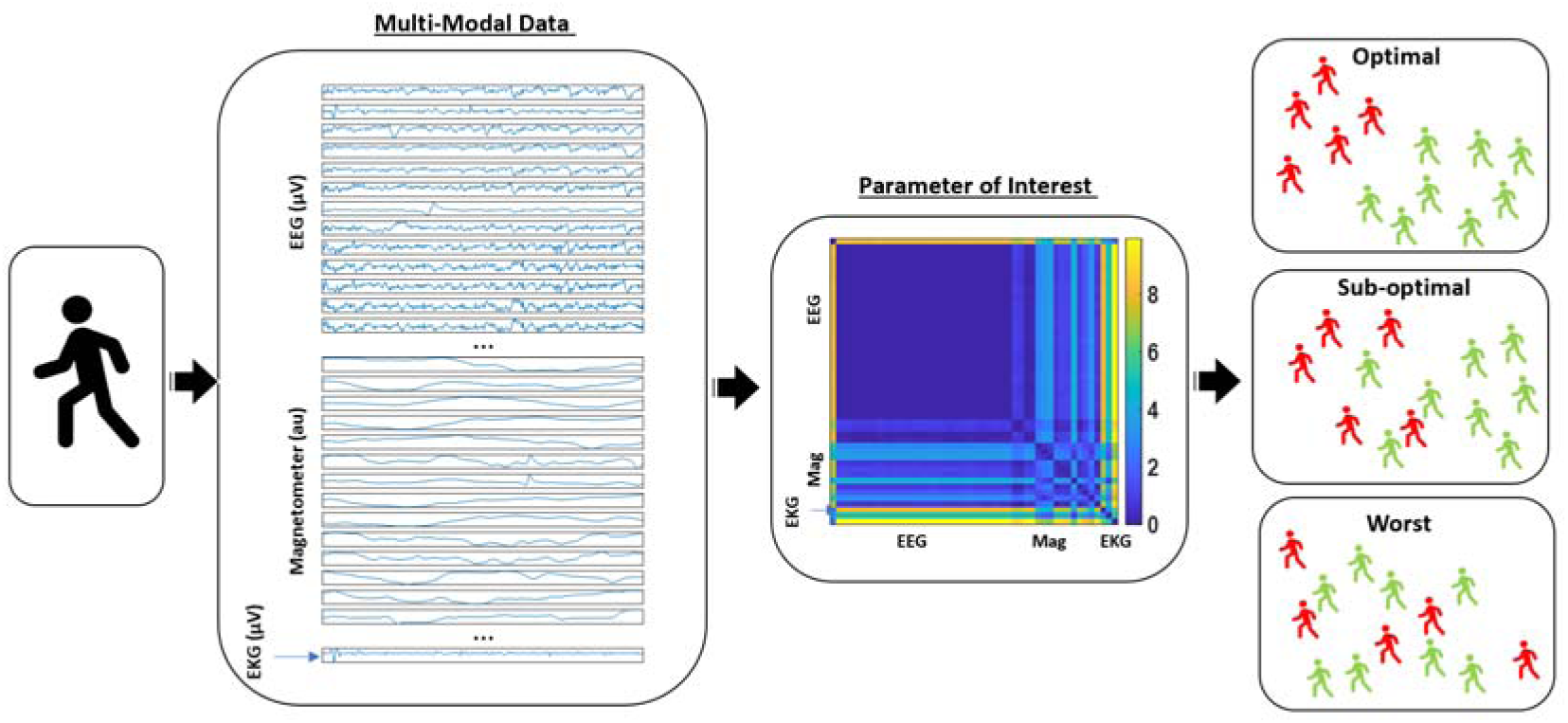
Multi-modal data of biophysical signals can be parameterized to characterize neurological features of PPD and identify optimal parameters, signals, and tasks maximally separating PPD from neurotypical aging. Sample continuous streams of data collected during a walking task using EEG, magnetometer and EKG data embedded in a wearable biosensor. With different sampling resolutions, different physical units, and altogether different patterns of individual variations in amplitudes and inter peak interval timings, several challenges lie ahead before we can make full use of digital data in clinical settings. One possible way to deal with such challenges is by identifying parameters in standardized form that span maximally separable dimensions and stratify cohorts into those with a disorder (such as PD) and those aging without neurological complications.

## 2. Materials and Methods

### 2.1. Participants

A total of 31 participants signed up and participated in this study. However, upon signal processing, we found that the data from 9 participants had too much instrumental noise or could not be properly synchronized across different devices for at least one condition. For that reason, the data from 22 participants is reported.

Among these 22 participants, 11 were healthy undergraduate students with ages ranging from 18 to 26 (10 female, 2 male). They were recruited from the Rutgers human subject pool system and received credit for their participation. The 7 PPD ranged in age from 64 to 77 years old (3 females, 4 males). They were recruited from the Robert Woodrow Johnson Medical Center at Rutgers University.

The Movement Disorders Society Unified Parkinson’s Disease Rating Scale (UPDRS) (*12*) for the PPD ranged from 16 to 44. The Hoehn and Yahr scale (*13*) ranged from 2 to 4. Additionally, 2 participants were healthy age-matched individuals with ages 65 and 68 years old (1 female, 1 male respectively). One was a spouse, and the other was recruited from ClinicalTrials.gov. Lastly, 1 participant was diagnosed with Essential Tremor (age 39, male) and 1 participant was diagnosed with ASD (age 15, female). These two participants did not show stark observable movement disorders, yet they were included as references too, because in any random sample arriving at a clinic, one could find such heterogeneous subtypes with invisible motor issues occurring at a microscopic level. These participants were recruited from ClinicalTrials.gov. The PPD, the patients with ET and ASD, and the aging controls received monetary compensation for their participation. Among the healthy young participants, one was left-handed, and among the PPD, one was left-handed. All participants had normal or corrected-to-normal vision. All participants provided informed consent, which was approved by the Rutgers University Institutional Review Board.

### 2.2 Instrumentation and Data Preprocessing

Participants wore two different types of wireless sensors - electroencephalography (EEG) and inertial measurement units (IMU) - to capture the biophysiological signals from the central (CNS), peripheral (PNS), and autonomic nervous system (ANS).

The wireless EEG device (Enobio; Barcelona, Spain) captured the brainwaves at 500Hz sampling rate with 31 sensors positioned across the scalp. The electrodes were spatially distributed as shown in Figure 2B, with the sensor Oz placed on the left abdominal region. This sensor was used as a proxy of electrocardiogram (EKG) to capture the heart signals. The wireless device was positioned on the back of the participant’s head, and the reference sensor was attached behind the left ear. Both EEG and EKG signals were recorded from the Neuroelectrics software (Enobio; Barcelona, Spain). Further preprocessing of the EEG signals were done in MATLAB-based toolbox EEGLab (*14*) and PrepPipeline (*15*) (MATLAB Release 2015b, The MathWorks, Inc., Natick, Massachusetts).

**Figure 2.**
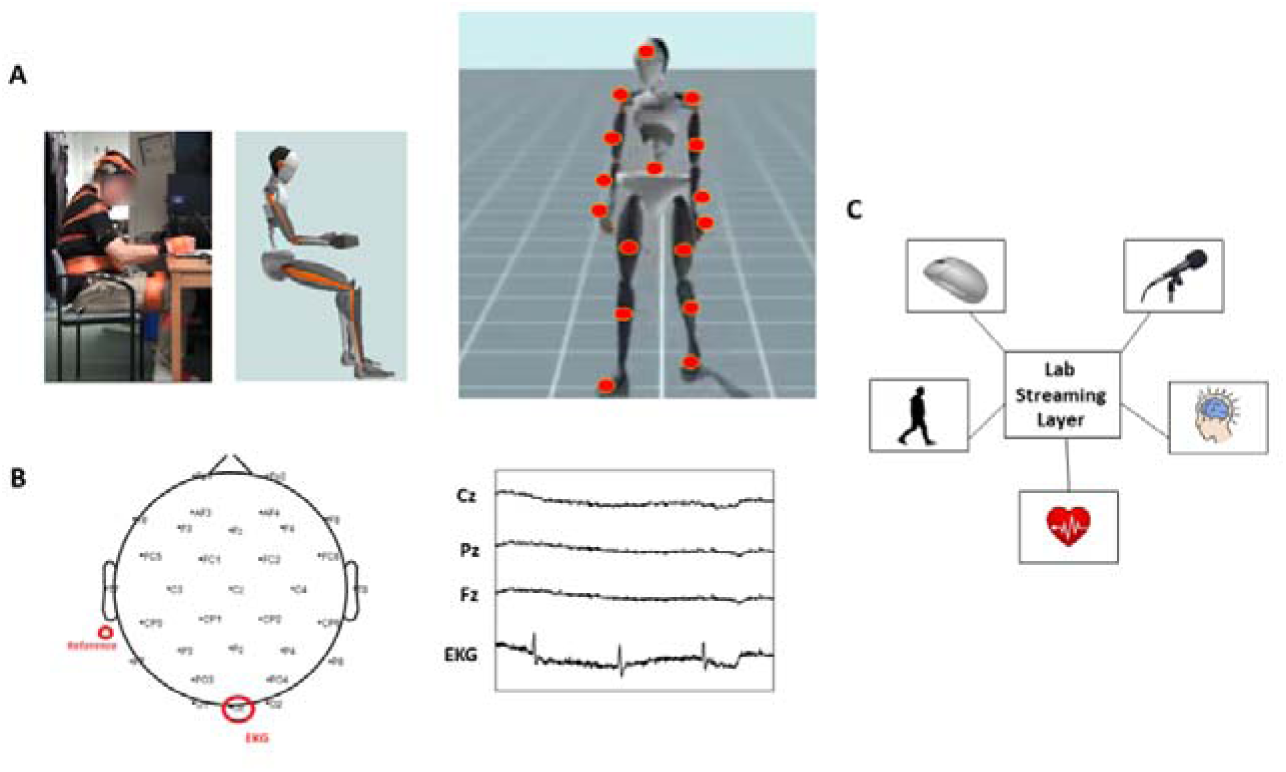
Instrumental Setup. (A) (Left) Picture of the participant performing a drawing task along with his rendered three-dimensional avatar registered in real-time. (Right) Bodily locations where we positioned the IMU sensors. (B) (Left) Location of EEG sensors, reference channel, and of the sensor (Oz) that recorded the heart activity; (Right) Sample EEG signals and EKG signal obtained from select EEG sensors. (C) Lab Streaming Layer synchronized the signals registered from multiple modes – mouse clicks, voice, kinematics, EKG, EEG.

Using the PrepPipeline toolbox, line noise at 60Hz was removed, and referenced via a robust average reference procedure, where channels were iteratively referenced to the average signal, while bad channels, such as those showing extreme amplitudes (deviation z-score exceeds 5) or lacked correlation with any other channel (correlation less than 0.4) were excluded and interpolated in this process. To eliminate the trend while preserving the cortical signals as much as possible, the EEG signals were further band-passed at 1-100Hz using Butterworth IIR filter at 1st order (*16*). The EKG signals were band-passed at 5-30Hz using Butterworth IIR filter at 1st order (*17, 18*) to eliminate trends while preserving the heart signal’s QRS complex as much as possible.

Note, because the heart signal was recorded with the reference channel positioned in less-than-optimal space (behind the left ear), the QRS complex were not clearly detectable with signal processing methods. This positioning of the reference was inevitable, because if we recorded the heart signal from a separate device and a separate software, with the reference channel positioned in the optimal location, there would be three different modes of signals recording through three different software’s on the same computer. This runs the risk of computer crash due to limited computation power. To avoid this risk, we recorded the heart signal by using one of the EEG sensors on the abdominal region and shared the same the reference channel with other EEG sensors registering the cortical signals. In this way, the computer co-registered signals from two devices (EEG and IMU) using 2 (instead of 3) software interfaces. This minimized the risk of computer overload and freeze, thus avoiding interruptions. However, the downside was the rampant noise in the obtained EKG signal and the inability to extract certain peaks within QRS complex. This tradeoff is important to consider when combining multiple wearables and synchronizing them to one CPU.

Motor signals from the PNS were captured with IMUs based on Xsens motion capture technology (*19*) sampled at 60Hz, and were acquired with Xsens MVN Studio software (Xsens; Netherlands). 17 sensors were attached to the participant with Velcro belts and additionally secured with athletic tape on the following body parts – head, sternum, posterior trunk, left and right shoulder, left and right upper arm, left and right wrist, left and right hand, left and right upper leg, left and right lower leg, left and right foot (Figure 2A right panel). These sensors allowed for creating the participant’s avatar (Figure 2A left panel), and registering each body part’s position, kinematics (linear speed (m/s) and angular speed (deg/s), linear acceleration (m/s^2^) and angular acceleration (deg/s^2^)), and magnetometer data (arbitrary unit; normalized from Gauss). For the purposes of this study, linear speed, position, and magnetometer data were mainly examined. Linear velocity, the first derivative of position, was chosen as it provided the least noise. From these velocity vector fields, the magnitude of the vector obtained at uniform time intervals (cm/s) was computed using the Euclidean distance. The magnetometer data was chosen for several analyses as it is relatable (and convertible) to the EEG signal, which is in units of μV.

In addition, the voice of the participant and the experimenter was recorded with a microphone sampled at 48,000Hz. However, the results from audio data are reported elsewhere (*20*).

These biophysical signals reflecting physiological states were temporally synchronized with the open-source package Lab Stream Layer (LSL). All software along with LSL were run on the same computer, and events were timestamped with mouse clicks on the display screen of that computer (Figure 2C). Because the EEG (cortical and heart) and the IMU (motor) signals were registered at different sampling rates, the EEG signals were down sampled to 60Hz. This ensured that all modes of signals were treated under a unified framework using a common data type.

### 2.3. Experimental Procedure

The setup of the instruments took about 30 to 45 minutes, which included donning the sensors and calibrating the systems. After the setup was complete, the participant performed a set of tasks that involved natural motions, which we digitized. These continuous streams of motions included drawing and walking. In this paper, we only present the results of the pointing and walking tasks further described below. The full description of the entire protocol can be found in (*21*), where different tasks and analyses are reported.

Both pointing (voluntary and requiring some planning) and walking (voluntary and highly automatic) tasks were used to probe different modes of action. These were designed to probe levels of spontaneous and deliberate entrainment of the biorhythms output by these biosensors and the external rhythm of a metronome. Both conditions were compared with a baseline mode, where no metronome was present. In each task, we used the following nomenclature - P1 and W1 represent baseline mode; P2 and W2 represent spontaneous metronome mode; P3 and W3 represent the deliberate metronome mode (explained below.)

#### *Baseline pointing without metronome* (P1)

The participants sat and pointed at a target with the dominant hand repeatedly 40 times at their own pace. For each pointing motion, the participant was instructed to start with the dominant hand in a resting position on the table, touch the target on the screen, and retract the hand back to its resting position.

#### *Spontaneous pointing with metronome* (P2)

The participants performed the same task as in condition P1 but did so while a metronome was beating in the background at 35 beats per minute. The metronome was set to play from a computer, and the volume was set approximately at 45 dBA. The participant was instructed to freely point at the target, but no instruction was given about the metronome. This ensured that any possible entrainment of the biorhythms to the metronome occurred automatically (rather than instructed.)

#### *Deliberate pointing with metronome* (P3)

The participants performed the same task as in P2 but did so under the instruction to pace the motions to the beat of the metronome (35bpm.)

This experimental assay was designed to probe differences in the entrainment of biophysical signals according to levels of mental intent (*i.e.,* deliberate intent *vs*. spontaneous self-emergent states) in relation to the hand moving at its own pace, without external metronome influences.

#### Walking Tasks

In the walking task, the participant freely walked around in an empty room for 15 minutes under the same three different conditions as above. In the baseline condition (W1), the participant naturally walked around the room at his/her own natural speed and in any direction for 5 minutes. In the second condition (spontaneous metronome W2), the participant walked for another 5 minutes but with the metronome beating in the background at 12 bpm. In the third condition (deliberately pacing the breathing to the metronome’s beat), the participant walked for another 5 minutes while instructed to pace the breathing rate to the metronome beat.

### 2.4. Data Analysis

In this section of the paper, a data transformation and four analytics will be discussed to capture the biophysical activities of the cohort while digitizing the bodily movements, heart, and brain responses to the tasks in the above-mentioned experimental assay. The goal here is to characterize the signatures of variability extracted from these waveforms and quantify the departure of these signatures in PPD from controls and from the two patients with other neurological disorders.

#### 2.4.1. Data Transformation: The Micro-Movement Spikes (MMS)

The position trajectories of the center of mass (COM) were registered with the Xsens MVN software (*22*) (Figure 3A.) We then obtained (by differentiation) the linear velocity vector field, and using the Euclidean distance, computed the magnitude of the vector per unit time, the linear speed (m/s) of the COM (Figure 3B). This was then land marked to automatically separate peaks and valleys, thus obtaining local maxima (red dots) and minima (black dots) respectively. The average amplitude of the time series was empirically obtained by fitting the best probability density function (PDF) and obtaining the absolute difference from the empirically estimated mean (based on the continuous Gamma distribution family). These absolute deviations were scaled using Equation 1, whereby the local peak amplitude was normalized by the sum of its value and the value obtained by averaging across points between the two surrounding local minima (see inset in Figure 3B). These spikes are referred to as the micro-movement spikes MMS) amplitude, and can be represented in a spike train form, as shown in Figure 3C.

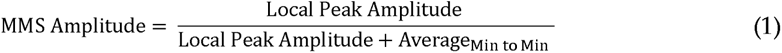

**Figure 3.**
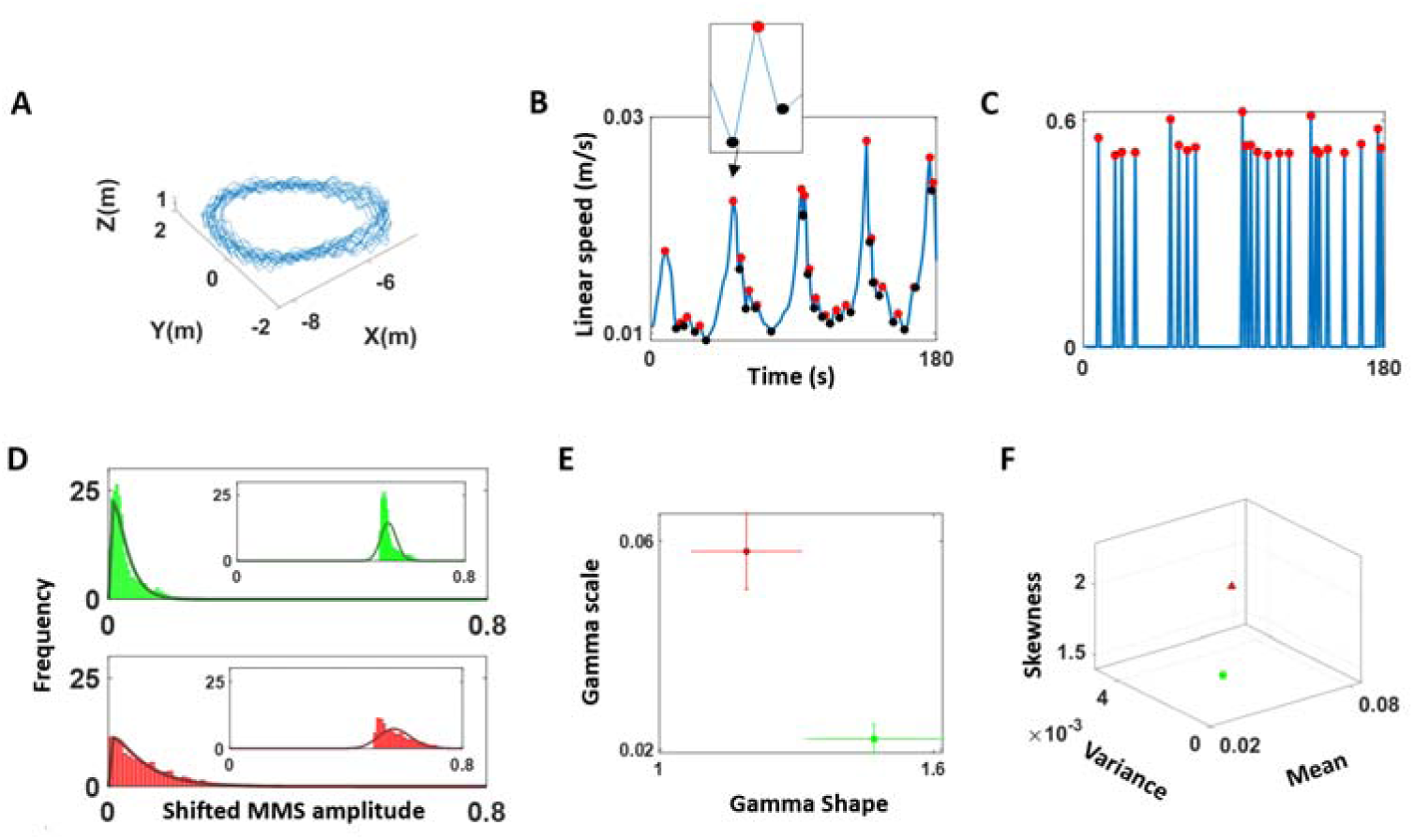
Pipeline of analyses of the COM speed timeseries. (A) The position of COM (m) was recorded at 60Hz. (B) Instantaneous linear speed profile was plotted as a waveform, and peaks (red) and valleys (black) of the linear speed (m/s) were extracted to compute MMS amplitudes using Equation 1. (C) The amplitude and of MMS peaks are represented in a spike train, where the values are 0 except instances of peak occurrences, which is set to MMS amplitude values. (D) Histogram of MMS amplitudes were plotted and found to have exponential fit (shown in insets). So, the values were shifted left by 0.5 to the left and were best fit in an MLE sense by the Gamma family (showing the Gamma PDF fit here). Top (green) and bottom (red) row show 2 different person’s data. (E) The best fitted Gamma parameters were plotted on a Gamma parameter plane with colors denoting different participants. (F) The first 3 moments of the fitted Gamma PDFs were computed and plotted on a 3-dimensional graph with colors denoting the different participants in panel F.

Such data transformation is commonly applied to avoid possible allometric effects due to individual anatomical differences (*23*). Furthermore, standardizing these moment-by-moment fluctuations in speed amplitude enables us to build unitless waveforms amenable to compare nervous systems readouts across different instruments with disparate physical units. We retain the original information about physical ranges while learning new features of the waveform variability that raises above local and global average states.

#### 2.4.2. Stochastic Analyses

The peaks in the MMS derived from the fluctuations in amplitude at the COM were plotted as histograms (insets in Figure 3D) where we found the density concentrated around 0.5, which is the average deviation in amplitude fluctuations captured by the MMS. We then shifted the entire time series by 0.5 and this allowed for a maximum likelihood estimation (MLE) of the best family of distributions to fit each of the histograms across conditions and participants. Figure 3D shows the best fit in an MLE sense, of the continuous family of Gamma PDFs. Figure 3E shows the Gamma parameter space spanned by the shape and scale values with 95% confidence intervals for two different sample participants. Figure 3F shows the parameter space spanned by the Gamma moments derived from the empirically estimated Gamma shape and Gamma scale parameters.

#### 2.4.3. Dynamical changes in relative distances to the COM

As a second set of analytics, we used the positions of all body parts and the COM at each time frame, and computed the distances of each body part in relation to the COM. Since walking is a cyclical task involving central pattern generators (CPG) (*3*) we can measure each body part, as it cycles relative to the COM. We can do so in three timesteps (t, t+1, t+2) and quantify how they collapse onto each other or remain independent from each other along the trajectory. To that end, we track these points unfolding on a three-dimensional parameter space spanned by the state of the fluctuations at each time frame. We measure the position of these frames along with the velocity and acceleration, relative to the vector (1,1,1). Then, the 3D datapoints are regressed to a plane for each body part (Figure 4C-E). In general, body parts that were the most active in its movement (such as the foot during walking, or the dominant hand during pointing) had the most circle-like shape with a hole in the center when projected onto the plane. These active cyclical body parts also had larger residuals of the plane fitting. (Details of the regression methods are described in Appendix C1.)

**Figure 4.**
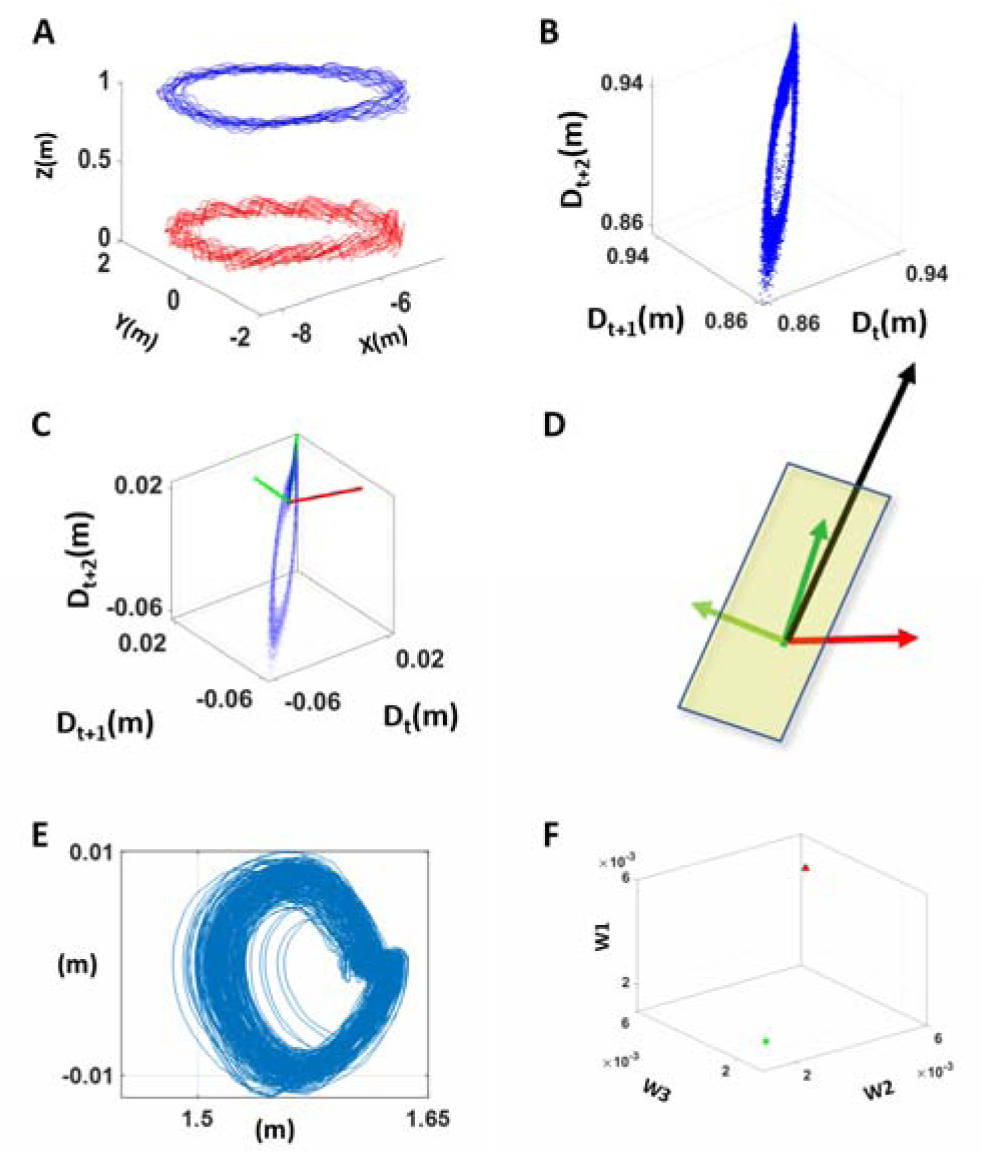
Analyzing the dynamical changes of body positions in relation to the COM. (A) Trajectory of positions of the right foot (red) and COM (blue) during walking. (B) Trajectory of distances between the right foot and the COM was represented in a 3-dimensional coordinate system, using the distance at times t, t+1, t+2 as the x, y, and z coordinate respectively. (C) The trajectory of the 3-dimensional coordinates was regressed onto a 2-dimensional plane, with the corresponding axes of the plane and the vector that is normal to that plane shown in green and red respectively. (D) The regressed plane is illustrated with unit vectors (axes) that reflect the most variation of the data marked in dark green, and the residual variation of the data marked in light green, and a vector normal to that plane marked in red. A normalized reference vector denoted in black is also shown to point towards the (1,1,1) coordinate, which represents the instance when there is no change in the distance between time t, t+1, and t+2. (E) The projection of the 3-dimensional coordinates in (B) on the regressed plane were plotted, and the shape of such projection was examined. (F) The parameter of interest was the deviation of the angle between the normal vector (red) and reference vector (black) in (D) from π/2 (*i.e*., abs(angle - π/2)). In general, a large deviation value would imply large motions across 3 consecutive time frames.

Given the 2 eigenvectors defining the regressed plane, and the normal vector of the fitted plane, we examined the angle between the normal vector and the reference vector. Here, the reference vector refers to the vector that spans (1,1,1) coordinate. If datapoints lie on this reference vector, we can interpret that there was no movement of that body part, since the distance is the same at time (frame) t, t+1, and t+2. The eigenvector with the largest eigenvalue is generally close to this reference vector, as the main datapoints hover around this reference vector, but deviates from it in relation to the magnitude of the body part movement. Hence, if the angle between the reference vector and the normal vector is exactly π/2, that implies that there was no movement. Conversely, if the angle deviates much from π/2, then that implies that there were large movements.

We analyzed this angle across the different tasks (P1, P2, P3) and (W1, W2, W3), and compared it between different cohorts of participants. Figure 4F provides an example with two participants localized as points on this (walking) parameter space. We underscore that incorporating the position of each body part at 3 sequential frames (time t, t+1, t+2) allows us to reflect important fluctuations in velocity and acceleration of the movement, relevant in characterizing PPD in relation to controls.

#### 2.4.3. Similarity Metric of Inter-Spike Intervals (ISI) across EEG independent components, EKG, and magnetometer data

To carry on the third level of analyses, we first extracted the ISIs from the MMS and then obtained the histograms. To compare the histograms across three different modes of data, we use the Earth Mover’s Distance (see Appendix C2) (*24, 25*), a distance metric to inform us about similarities and differences of relevance across participants and conditions.

We used three modes of data to obtain the ISIs – EEG to reflect the CNS, magnetometer to reflect the PNS, and EKG to reflect the ANS. To represent the PNS, we chose to use the magnetometer data, since its unit is a function of voltage comparable to EEG and EKG units. The motion capture device used in this study yields magnetometer data in arbitrary unit (au), which is a normalized value of Gauss unit. Because we focused on the timing of the spikes, and not the amplitude, the normalization of the values should not affect the results of this analysis.

The EEG data were decomposed through independent component analysis (ICA) using the Infomax ICA algorithm (*14, 26, 27*) with at most 512 iterations. Such decomposition allows locating a set of sources from which the signal originated from. This decomposition was done per task (condition) and provided a set of time series of 15 to 31 components (depending on how many rank/null spaces were present across task/participant), which were a set of spatially static, and maximally independent component processes. Subsequently, source localization of each of these components was performed, using Dipole fitting method (*28*) that co-registered to fit the scalp topography of a Spherical Four-Shell (BESA) head model. The locations of the obtained sources of each independent component were then categorized into ‘in-brain’ and ‘out-brain’. Components that were categorized as ‘in-brain’ were those with sources located within 83 mm from the center of the head model; and sources located beyond 83 mm from the center of the head model were categorized as ‘out-brain’. Given that channel sensors were located uniformly 85 mm away from the center, 83 was a conservative choice that accounted for sources that were located on the scalp, presumably reflecting the head and facial muscle movements. The separation of components inside and outside the brain allowed us to distinguish signals that are generated within the brain from those generated from the muscle and other non-brain related artifacts. In general, there were approximately 0-5 components (out of 15-31 components) whose dipole locations were categorized to be ‘in-brain’, and the rest were considered ‘out-brain’. In our study, the residual variance of a typical fitted dipole during the walking tasks ranged from 5% to 45%. Given the small number of in-brain components per task/participant, excluding some components with large residual would lead to loss of data altogether. For that reason, we kept all components that were provided by the dipole fitting, while taking notice of the diverse range of model fitness.

After the ISIs of the EEG components, magnetometer, and EKG were gathered for each condition per participant (Figure 5A), histograms of ISIs were created for all EEG sensors, all magnetometer sensors, and EKG sensor, yielding 33-49 histograms per condition (depending on how many independent components were derived from the EEG data) (Figure 5B). These histograms were constructed as single probability distributions with bin width as 1 frame (1/60s), minimum bin value as 1 (frame), and maximum bin value as the lower of 30 and the maximum ISI. Any ISI that exceeded 30 frames (0.5s) was not included in this analysis, as this would most likely be due to instrumental noise. Based on these histograms, pairwise earth mover’s distance (EMD) was computed to understand the shifts in stochasticity of signals across different tasks. Details of the EMD algorithm is elaborated in Appendix C2. Finally, the median EMD values were compared per different pairs of sensor categories (*e.g.,* pairs comprised of 2 EEG sensors; pairs comprised of 1 EEG sensor and 1 body part’s magnetometer; pairs comprised of 1 EKG sensor and 1 body part’s magnetometer), and across the three conditions (Figure 5C). To compare across cohorts, we plotted the median EMD values from the three conditions as a single datapoint for each participant with different colors per cohort, to observe any clustering patterns that would emerge.

**Figure 5.**
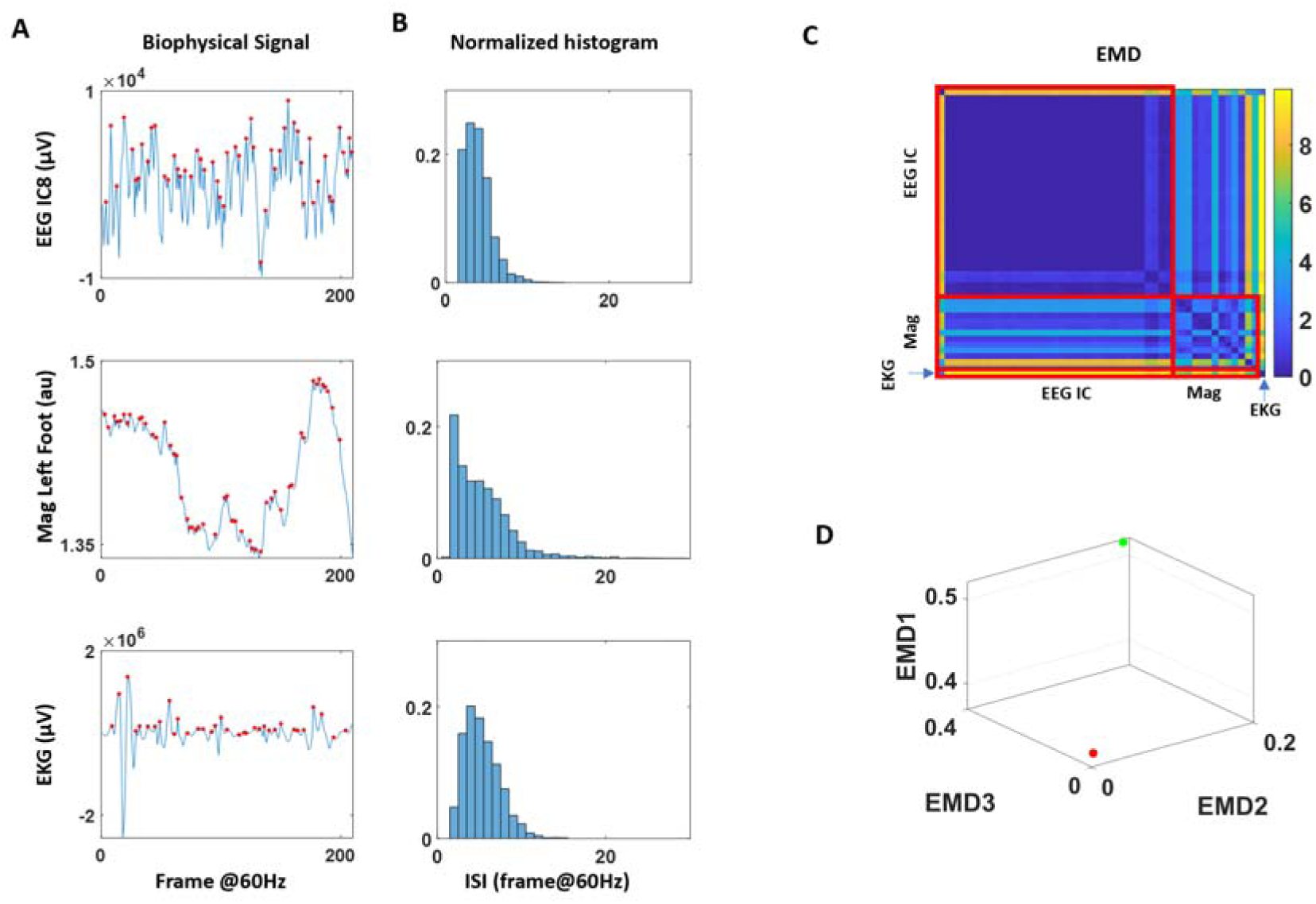
Analyzing EMD of inter-spike intervals (ISI) across 3 modes of signals. (A) Biophysical waveforms of the EEG independent component in μV (top), magnetometer in arbitrary unit (middle), and EKG in μV with spikes marked in red. (B) Inter-spike interval (ISI) times were computed for all sensors, and a frequency histogram was plotted. To make these histograms comparable across different modes of sensors, the frequency was normalized to values between 0 and 1. (C) Matrix of EMD for all pairs of sensors was plotted for each participant/task. (D) For each category of sensor pairs, the median EMD was computed for each condition. The median EMD during the control condition (P1, W1) was represented on the z-axis (EMD1), the metronome spontaneous condition (P2, W2) on the x-axis (EMD2), and the metronome with deliberate, paced condition (P3, W3) on the y-axis (EMD3). Two sample participants from different demographic in the cohort are localized in this parameter space and differentiated by color.

To understand the relations of EEG sensor signals with magnetometer/EKG signals, with respect to those generated from within the brain and outside the brain, the same analysis was done by further subdividing the EEG component by their dipole locations, as either ‘in-brain’ or ‘out-brain’. While the above analytics examine the overall EMD values across different participants, we also examined the variability of the EMD values between different sensor pairs. Specifically, for each participant, there are approximately 800 EMD values per condition, since there are approximately 40 ISI histograms per participant (ranging between 33 and 49). As such, a pairwise computation of EMD on these ISI histograms would yield approximately 800 MI values. Based on these EMD values, we compiled a set of 3 dimensional coordinates, by taking each pair of sensors, and plotting their EMD values at control (P1 or W1) on the z-axis, metronome spontaneous (P2 or W2) on the x-axis, and metronome deliberately paced condition (P3 or W3) on the y-axis (Figure 6A).

**Figure 6.**
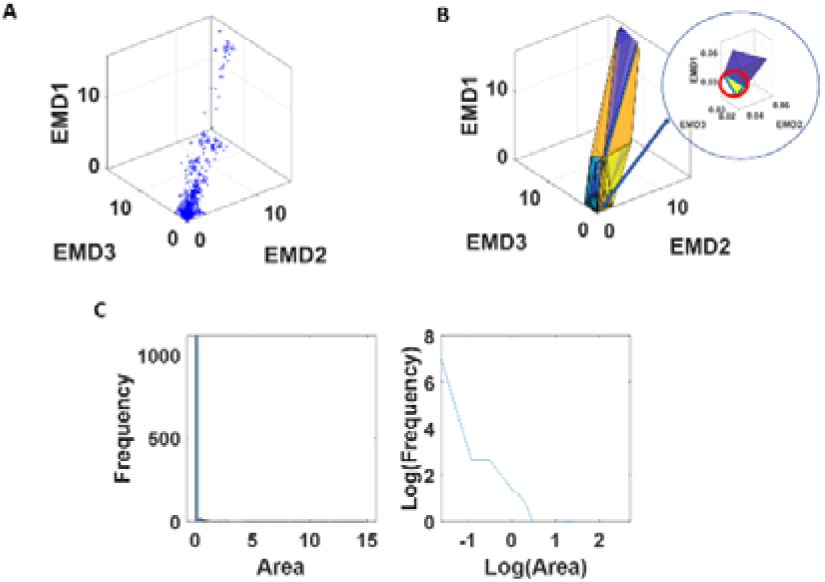
Analyzing variability of dynamics characterized by EMD. (A) For all sensor pairs, the corresponding EMD values from the control condition (z-axis, EMD1), spontaneous metronome condition (y-axis, EMD2), and deliberately paced metronome condition (x-axis, EMD3) were plotted. (B) Delaunay’s triangulation revealed triangles across the 3-dimensional coordinates, and the areas of these triangles were computed. (C) Frequency distribution of the linear and of the log-log values, whereby slope and intercept of the latter were obtained and further analyzed.

Note, the number of total sensors is different for each condition, as the number of EEG IC are different across tasks. For that reason, when comparing the change across conditions for pairs that included the EEG IC, the choice of EEG component was random. For example, if there were a total of 10 EEG components in the control condition, 12 components in the metronome, and 14 components in the paced condition, we took the minimum number of components (*i.e.,* 10) from the 3 conditions, and randomly chose 10 components among the 12 in the metronome, and 10 components among the 14 in the paced condition. As a result, the 3-dimensional coordinates pertaining to the EEG data do not exactly correspond to each other. That is, the x- and y- coordinate of the paired sensors that include the EEG IC do not exactly correspond to the EMD of the pairs represented in the z-coordinate. Nevertheless, given the large amount of data used in these analytics (∼800 datapoints per person), characterizing the patterns of different participants, such differences are negligible.

Once these 3-dimensional coordinates were obtained, the scatter of points formed surfaces that we characterized using the Delaunay triangulation. Given a set of discrete points, the Delaunay triangulation (*29*) creates a matrix of N x 3, where the 3 columns represent the datapoint indexes that form the 3 vertices of each N triangle. These triangles are formed such that no other point is inside the circumcircle of the formed triangle (Figure 6B). Here, we took a list of these such triangles and computed the surface area for each triangle. We then plotted a frequency histogram of these points (Figure 6C). The triangle areas represent the spread of these datapoints, whereby a larger value would imply a wider range of EMD (larger shifts in stochasticity) across conditions.

Because of the large number of triangles and skewed distribution of such histograms (as shown in Figure 6C - left), these were examined by applying logarithm to both axes of the histogram (Figure 6C - right). Given the linear shape of such power-law distribution, we then regressed this to a line, to examine the spread of such triangle areas. Essentially, the wide spread of triangle areas (represented by a flatter slope and lower intercept of the power-law distribution) would imply a wider range of EMD between different sensors; conversely, a narrower spread of triangle areas (represented by steeper slope and higher intercept of the power-law distribution) would imply a narrower range of stochasticity between different sensors.

This way of representing the EMD data is different from those shown in Figure 5D, which essentially reflects the median triangle areas; that is, the median change in stochasticity across conditions. Here in Figure 6C, the shape of the power-law distribution of triangle areas reflects the *variability* of stochastic shifts between different modes and location of sensors.

#### 2.4.4. Pairwise Cross-Correlation of Magnetometer Data

As a last set of analytics, cross-correlation was examined between different pairs of magnetometer sensors (Figure 7A). Here, the time series of magnetometer data were normalized by linearly scaling the values to range between 0 and 1 (Figure 7B). Then, they were separated into 5-second window segments. For each segment, cross-correlation was computed for each sensor pair (Figure 7D), and the median of these cross-correlation values for all pairs obtained from each window. The median values, taken across all time window segment’s medians (of all paired sensors) were computed to provide a single summarizing cross-correlation value per condition. These were then plotted for all participants, whereby the z-axis represented the median cross-correlation value during the control condition (W1 or P1), and x- and y-axis used to represent the values during the spontaneous metronome (W2 or P2) and deliberately paced metronome (W3 or P3) condition respectively (Figure 7E).

**Figure 7.**
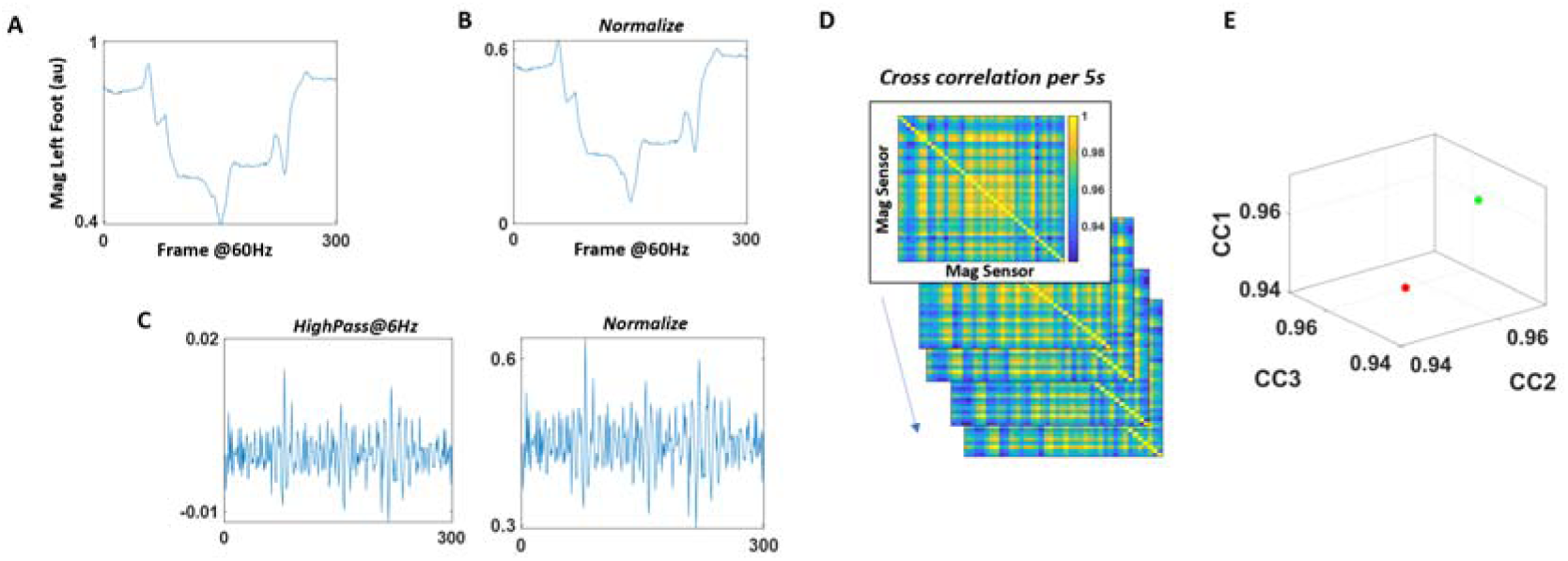
Analyzing cross-correlations between different magnetometer sensor data. (A) Magnetometer data of the left foot in arbitrary unit is plotted. (B) For all body sensors, the raw data was linearly scaled and normalized to range between 0 and 1. (C) For all body sensors, the raw data was also high-passed at 6Hz (left), and then normalized to range between 0 and 1. (D) For each 5 second segment, cross-correlation was computed between all sensor pairs for both normalized magnetometer data (from (B)), and high-passed and normalized magnetometer data (from (C)). Pairwise maximal cross-correlation value was extracted and plotted on a matrix. (E) The median maximal cross-correlation was computed for all sensor pairs and for all 5-second window segments, and was plotted on a 3-dimensional plot, where the z-axis (CC1) denoted the median value during control condition (P1, W1), x-axis (CC2) denoted the value from the spontaneous metronome condition (P2, W2), and y-axis (CC3) denoted the value from the deliberately paced metronome condition (P3, W3). The marker color denoted the different participant type in the cohort.

We also computed cross-correlations of magnetometer data that were high-bandpassed at 6Hz using Butterworth IIR filter at 2nd order and compared these across the different participants. Tremor is one of the main symptoms of PPD. This tremor of the body motion is known to exist in the 4-6Hz range (*30–32*). For that reason, we excluded the tremor by high-pass filtering the magnetometer data, to understand how the tremor impacts characterizations based on the cross-correlation analytics (Figure 7C).

Note, the same analysis was done on the EEG component data, and similar results were found as with those from using magnetometer data. However, we found the magnetometer data to be more informative in characterizing PD than the EEG data. Hence, for simplicity, in this study, we focus on the results from the magnetometer data.

## 3. Results

### 3.1. Stochastic Signatures: Higher NSR of COM Revealed in Walking Task

The trajectory of the COM was first examined for each participant during their walking tasks. When we compare the baseline condition (W1) across all participants, there is a noticeable difference in its regularity. The neurotypical (NT) participants show a pattern of regular cycles in their walking trajectories. In contrast, PPD show more variable and degraded cycles, as shown in Figure 8A. To quantify such pattern, the MMS derived from the fluctuations in speed amplitude were best fitted in an MLE sense, by the Gamma PDF (with 95% confidence intervals for the shape and scale parameters.) This was amenable to interpretation, allowing us to understand the stochastic nature of the COM kinematics.

**Figure 8.**
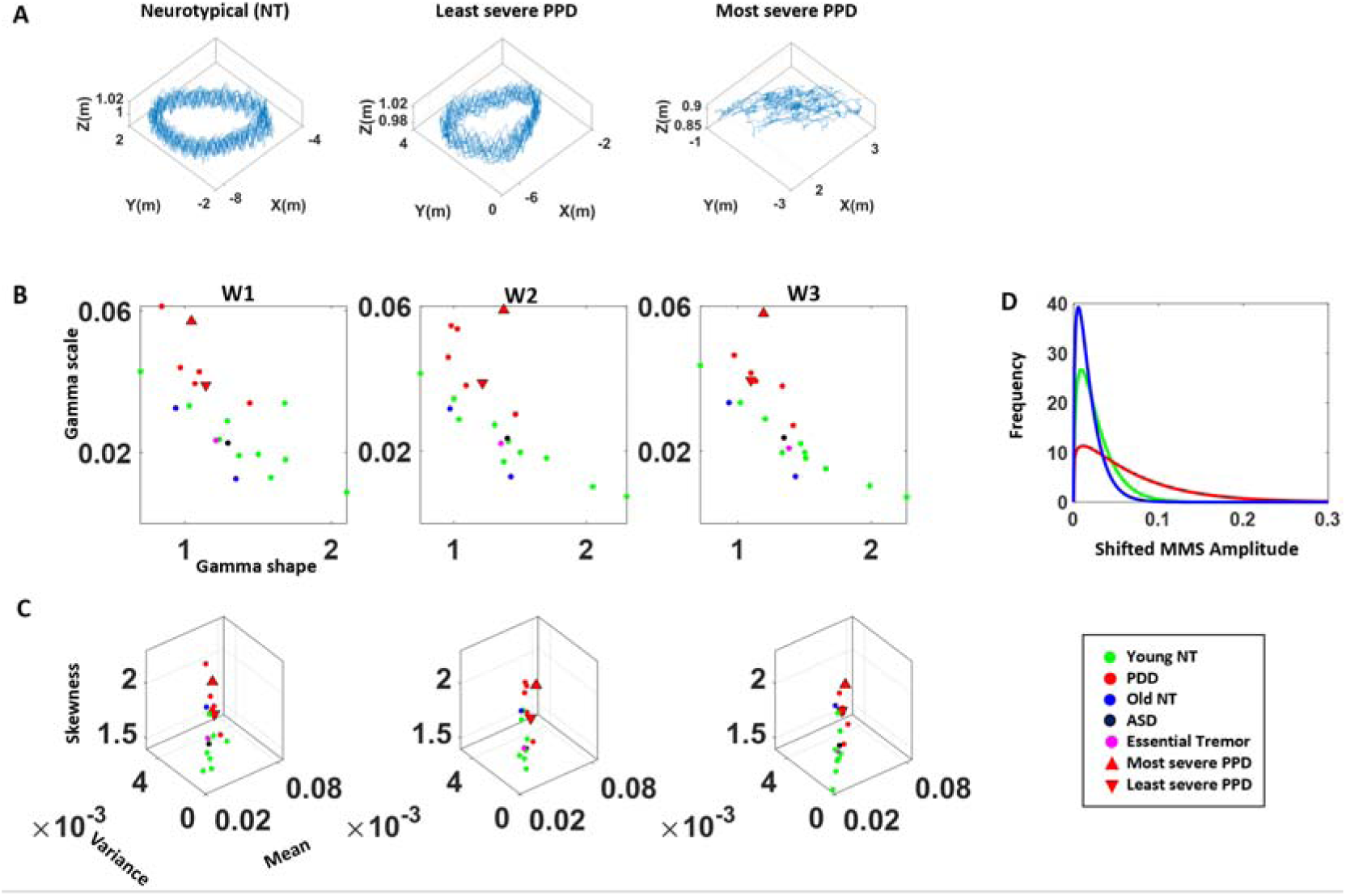
Clinically interpretable stochastic signatures of the COM during walking tasks. (A) Representative COM trajectory during a walking task (left) of a healthy young (NT) participant, (middle) of the PPD with lowest severity, and (right) of the PPD with highest severity (according to clinical criteria.) (B) Fitted Gamma parameters (MMS amplitudes from COM linear speed) are plotted on the Gamma parameter plane for each walking tasks (condition W1, W2, W3), with marker color differentiating participant type. (C) Corresponding Gamma moments of the fitted PDF (x-axis denotes mean; y-axis denotes variance; and z-axis denotes skewness) for each walking task of the assay. (D) Fitted PDF of frequency histograms (of shifted MMS amplitudes from the COM linear speed) of representatives NT participant (green), healthy age-matched participant (blue), and PPD (red).

When examining the fitted Gamma parameters, consistent with the patterns found in previous studies on PPD (*33*), we generally found higher values of the Gamma scale and lower values of the shape parameter in the patients, with stochastic shifts across conditions. These departed from NT controls across all three walking tasks (Figure 8B). Noticeably, the most severe patient (according to the clinical scales) is localized at the extreme (highest scale value, which is the highest NSR). The least severe patient localizes at the boundary of the cluster between the NT and the PPDs. When we examine the three Gamma moments, skewness is noticeably high for the PPD, compared to their NT cohort. In correspondence with the Gamma parameter plane, the most severe PPD is localized at the highest skewness level, relative to the rest of the participants. Alongside, PPDs show a higher mean and variance shown by its flatter distribution in its fitted PDF. This is contrasted with their NT cohort in Figure 8C, 8D.

In addition to these, one age-matched (elderly) NT participant shows a slight difference from the younger NT cohort, as its fitted parameters and moments are positioned between the clusters of PPD and the younger NT cohort. The other age-matched NT does not show much difference in relation to the younger cohort. Likewise, the patients with ASD and ET do not show much difference from the NT cohort.

### 3.2. Reduced Dynamical Changes in Relative Body Parts’ Distances to the COM Characterize PPD

The trajectory of the distance between right foot and COM were plotted on 3-dimensional graph with coordinates as the relative distance at time frames (t, t+1, t+2). These datapoints were then regressed onto a plane, and the projection of these is plotted in Figure 9. Here, we find contrasting shapes in the projection between the NT cohorts and the patient cohorts. The NT projections form a donut-like shape with a hole in the middle. In contrast, patients (Parkinson’s, ASD, ET) do not show such shape. Instead, they show a rather squashed version.

**Figure 9.**
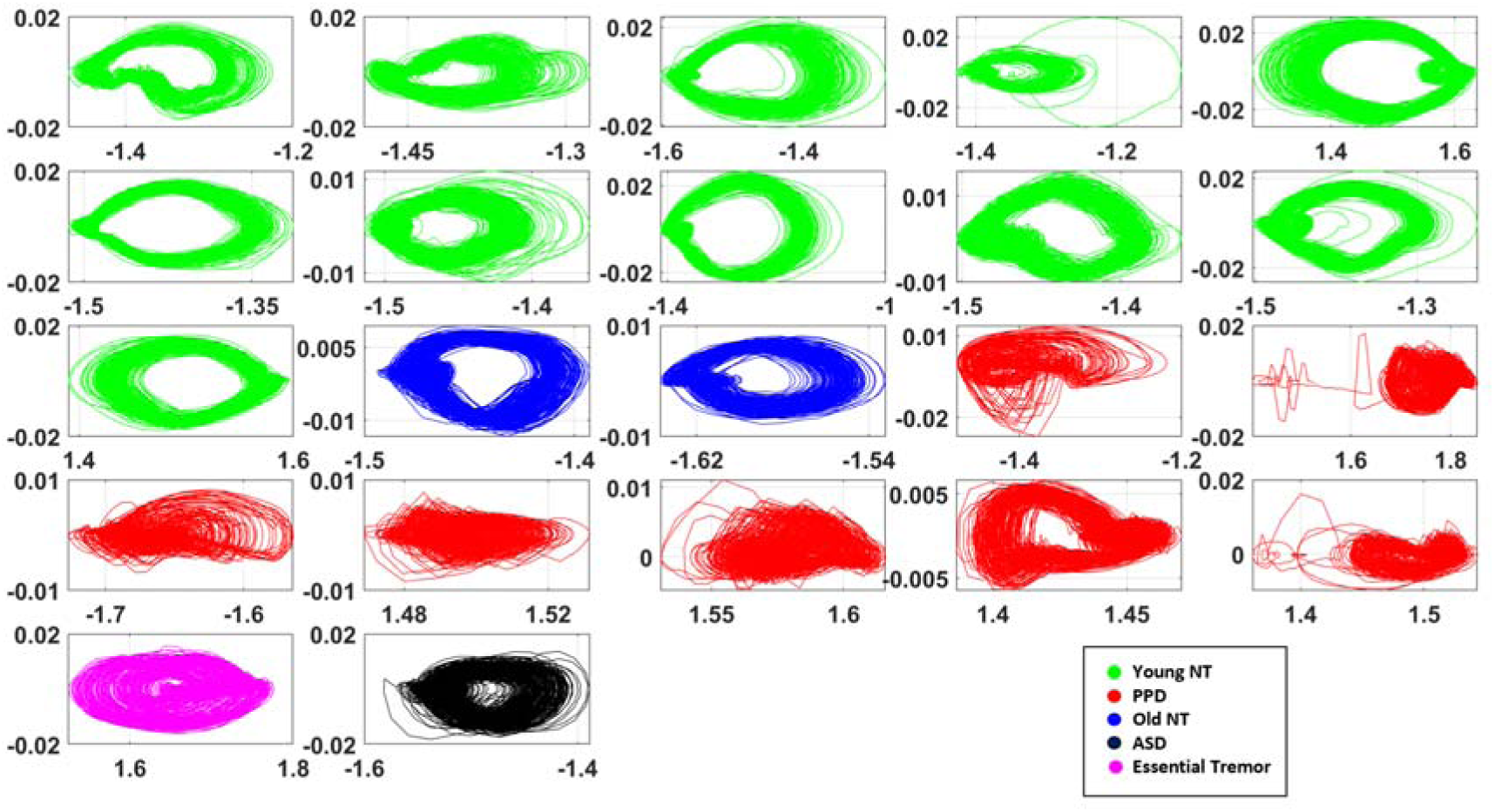
Projection of the 3-dimensional coordinates (of distance between right foot and COM). For most Parkinson’s patients (red), we observe an absence of a donut-like shape in the projected trajectories, contrasted by the NT participants (green, blue).

The hole in the middle reflects the distance in datapoints diverging from the reference vector (1,1,1). If the body part moves fast at each moment, the instantaneous frame-by-frame change in position of the body part is large. This is most often the case for active body parts (*i.e.,* foot). In such a case, the datapoints tend to diverge from the reference vector. If this fast action is regular and cyclical, the projection trajectory exhibits a hole in the middle. In contrast, when the motion is irregular and a cyclical (*e.g.,* due to loss of balance and on-line corrections), the hole in the middle disappears. The NT controls show the healthy pattern, whereas the patient participants tend to have a slower and irregular pattern in their trajectory cycle.

To quantify the extent to which these distance datapoints deviate from the reference vector, the angle between the reference vector and the normal vector of the regressed plane were computed. If the datapoints did not deviate at all (*i.e.,* there were no motions), this angle would be exactly π/2. Conversely, if the datapoints have large deviations (*i.e.,* there was a persistence of cyclical fast motions) this angle would largely depart from π/2. This departure was plotted for all participants for the left toe in Figure 10A (left), with each axis denoting different conditions of the walking task (W1, W2, W3). Overall, the clustering pattern is consistent across all body parts as shown in Figure A1, such that NT cohorts tend to have a large angle deviation than the PPD, and this is the case for all three conditions. Walking segregates the PPD from the NTs with high statistical significance across all conditions.

**Figure 10.**
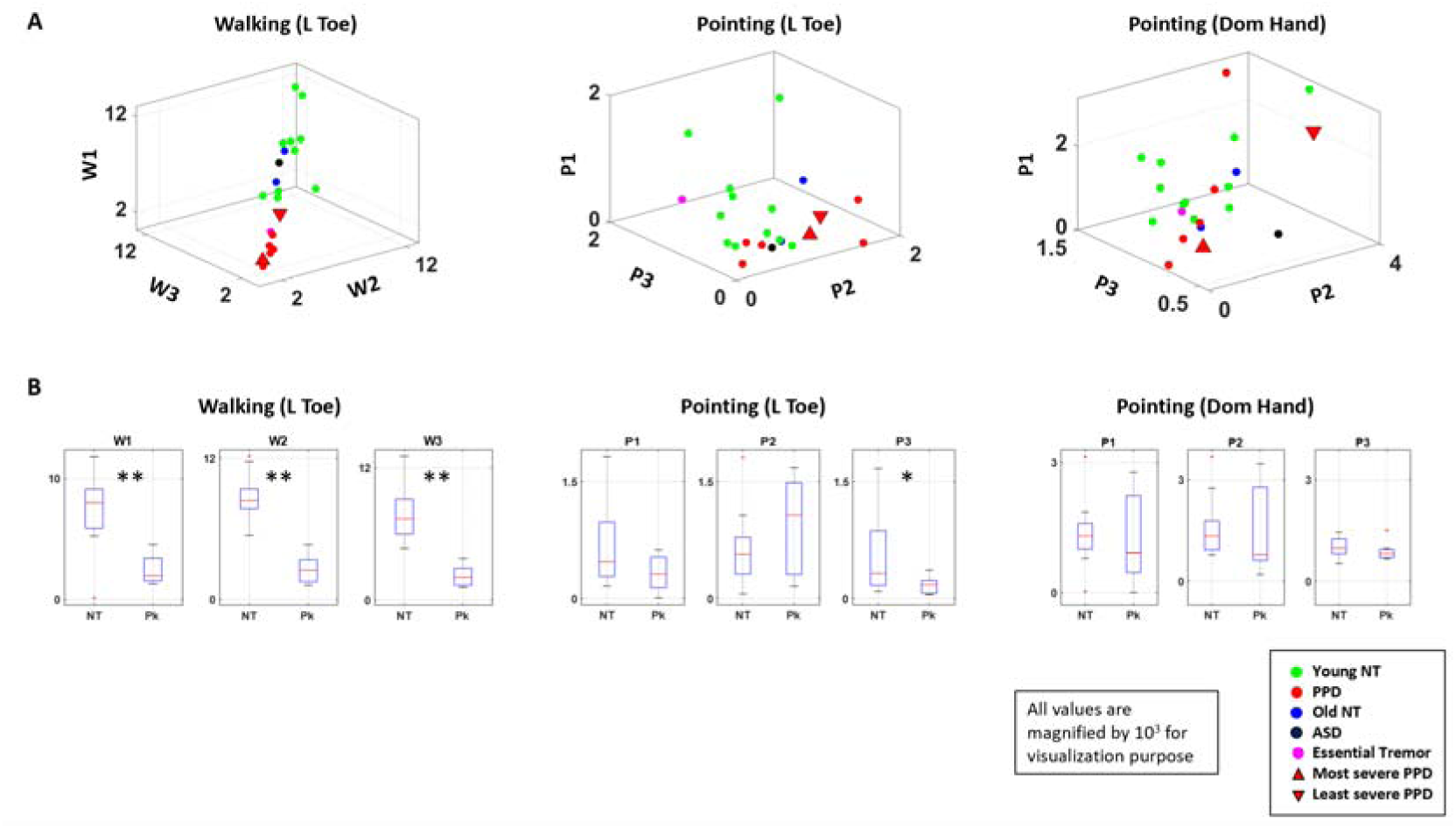
Interpretable digital biomarkers and identification of task and parameters to differentiate PPD from NT. (A) Angle deviations from π/2 of walking *vs.* pointing. (Left) Angle deviation of the left toe during walking tasks (W1, W2, W3), (middle) left toe during pointing tasks (P1, P2, P3), (right) and dominant hand during pointing tasks (P1, P2, P3) are plotted for each participant with color denoting the demographic cohort. (B) Box plot of the angle deviation values for NT and PPD cohort for all 3 conditions of walking and pointing tasks, with statistical significance denoted by * (p<0.05) and ** (p<0.01).

The two age-matched (elderly) NT participants generally lied within the cluster of the younger NT cohort, but close to the boundary between the NT and Parkinson cluster. The ASD patient’s datapoint generally lied within the cluster of the younger NT cohort, while the ET patient’s datapoint is localized within the PPD cluster.

#### Parameter Identification

For further comparison, we took the distance between the left toe and the COM as our parameter of interest for walking tasks, and the distance between the dominant hand and the COM as the parameter of interest for the pointing task. These body parts were chosen, because the most active body part revealed the highest separation between different cohorts. We also examined the distance between the left toe and the COM during pointing tasks, to compare outcomes between the walking and the pointing tasks (Figure 10A).

During the three pointing tasks, for the left toe, there is some difference in clustering pattern across the different tasks. Specifically, there is little separation between the PPD and the NT participants for conditions P1 and P2. However, there is statistically significant separation between the two cohorts for condition P3, such that the PPD tend to have little angle deviation, while the NT participants tend to have larger angle deviation. Among the two age-matched NT participants, one’s datapoint lied closer to the cluster of PPD, while the other participant’s datapoint lied closer to the NT cluster. Also, the ET participant is localized closer to the NT cluster, while the ASD participant, is localized closer to the cluster of PPD. This was also shown as a trend for the dominant hand during the pointing tasks (although with no statistical significance). There we found similar clustering patterns, with more separation between the clusters of NT and PPD in condition P3 (Figure 10B). See Table B1 for details of the statistics.

In general, patients with impeded motor control seemed to move slower, which is well captured by the lower angle deviations. However, this was only noticeable during the walking tasks. This was not necessarily the case for pointing tasks, implying that in PPD the walking task (requiring balancing the full body) is highly revealing of the motor control problems across different modalities and levels of intent. This walking task was very revealing (with high statistical significance) of differentiations (or lack thereof) between deliberate and spontaneous levels of entrainment with the external rhythms of the metronome.

### 3.2. Smaller and more variable range of EMD between EEG independent components, EKG, and magnetometer data among PPD

The median EMD of ISI histograms between different modes of sensors were examined during the 3 walking tasks for all participants and plotted for all participants in Figure 11. We examined the interactions by different categories of sensor pairs. For all categories, PPD tend to have lower EMD values than the NT cohorts for all sensor pairs. The difference was found statistically significant for the pairs of 2 EEG component data, 1 magnetometer data and 1 EEG component data; 2 magnetometer data, 1 EKG and 1 magnetometer data. Details of the statistical test can be found in Table B2.

**Figure 11.**
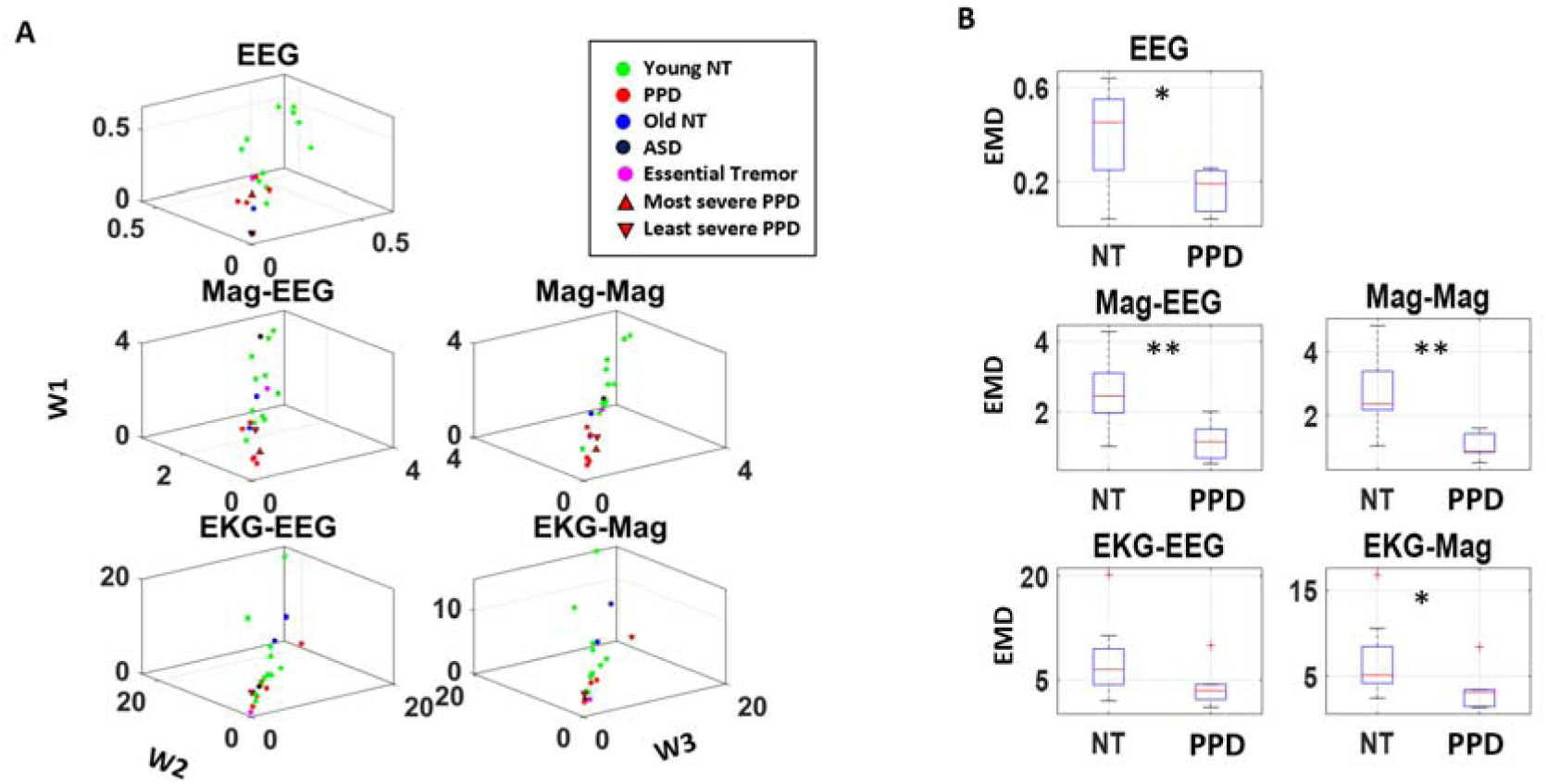
Identification of biophysical signals with highest change in stochastic signatures, using the EMD of inter-spike interval (ISI.) (A) Median EMD for condition W1 is plotted on the z-axis (W1), for condition W2 on the x-axis, and for condition W3 on the y-axis; for different sensor pair categories (reflected in each subplot) of each participant. Colors in legend denote different demographic cohort. (B) Box plot of median EMD values were plotted and compared between NT participants (including young NT and age-matched NT) and PPD. For all modes of signals, PPD show a lower EMD, identifying the magnetometer data as the most evident.

To see if such pattern persists when the EEG component data is analyzed separately for those that are generated by the brain activity, and those that are generated by non-brain activity (*e.g*., muscle motion), the same analysis was done as is shown in Figure 12. There we show EEG component data from inside (IN EEG) and outside the brain (OUT EEG).

**Figure 12.**
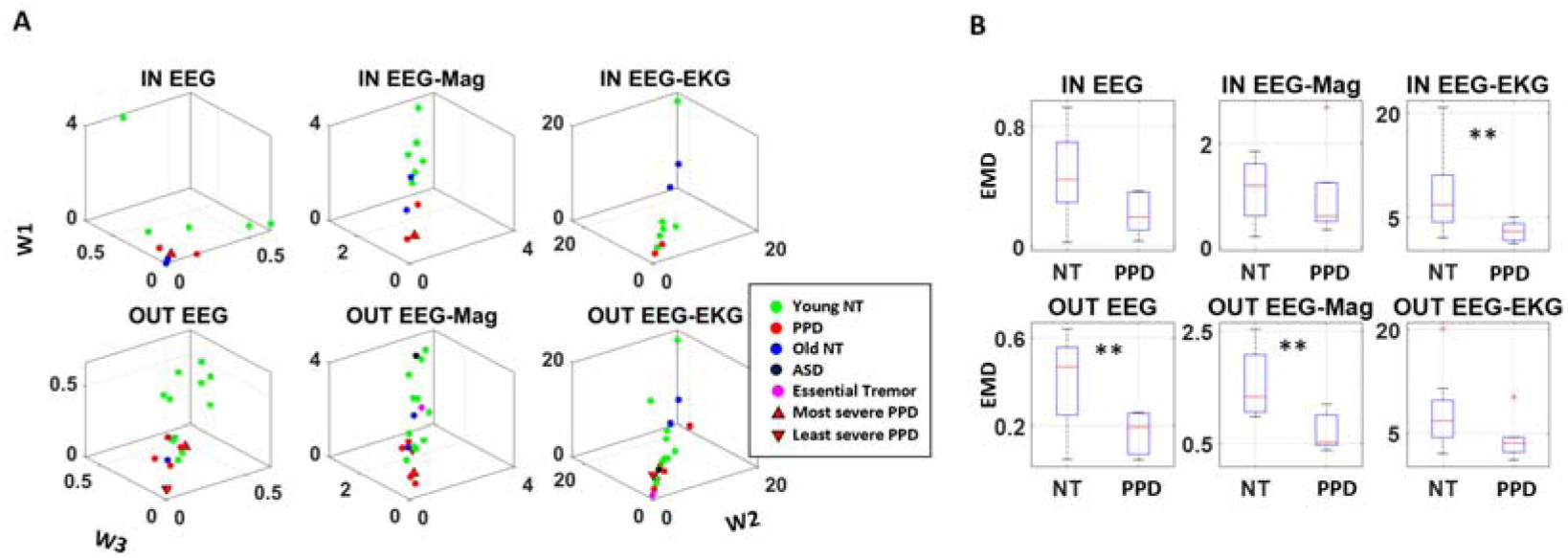
EMD of inter-spike interval (ISI) for sensor pairs with EEG IC localized inside vs. outside. (A) Median EMD for condition W1 is plotted on the z-axis, for condition W2 on the x-axis, and for condition W3 on the y-axis; for different sensor pair categories (subplot) of each participant, with colors denoting different demographic cohort. Top row shows the sensor pairs that include EEG IC localized within the brain, and bottom row shows the sensor pairs that include EEG IC localized outside the brain. (B) Box plot of median EMD values were plotted and compared between NT participants (including young NT and age-matched NT) and PPD for sensor pairs that include the EEG IC data localized within the brain (top) and outside the brain (bottom).

Although the pattern remains the same, whereby the overall EMD is lower for the PPD than the NT cohort, the statistical significance is pronounced in the pairs comprised of 1 OUT EEG and 1 magnetometer data, 2 OUT EEG data, and 1 IN EEG and 1 EKG data. Given that much of the EEG component localized outside the brain are mostly due to motion artifacts (*14*), it can be assumed that signals coming from the PNS tend to drive the changes in stochastic signatures. However, we also notice a strong pattern between IN EEG and EKG which begs the question of whether ANS in the PNS drives the change, or whether the change is centrally driven. (See further statistical test results in Table B3).

We further analyzed the overall variability of EMD values across different modes and spatial location of sensor pairs within each condition. To that end, we plotted the histogram of the area of the estimated Delaunay triangles spanned by the surface of the scatter all sensor pair datapoints. The log-log transformation of this histogram was then used to fit a line. The slope of this regressed line was the parameters of interest. Here a steeper slope implies less variability of EMD values across sensor pairs. Conversely, a flatter slope implies more variability of EMD values.

The PPD generally show a flatter slope than the NT cohort. This hints at more variability across their stochasticity, while exhibiting an overall lower EMD value shown by the lower intercept (Figure 13). In this context, the typical PPD shows a wider variability of the EMD values, implying that their stochasticity is more variable across context (condition) and type of signals (sensor).

**Figure 13.**
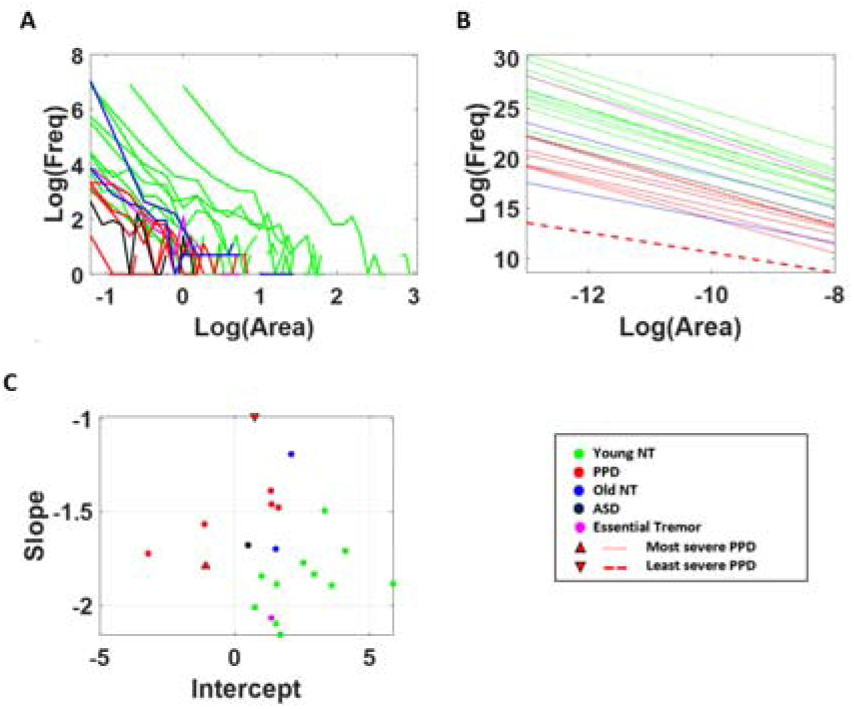
Capturing variations in stochastic shifts in PPD. (A) Triangle areas formed by Delaunay’s triangulation method of EMD datapoints of all sensor pairs were computed, and its frequency histogram obtained as a log-log plot, with colors denoting different demographic cohorts. PPD tend to have smaller area value, implying lower level of stochastic shifts. (B) The log-log plot of the frequency histogram is linearly regressed for each participant. PPD have a lower intercept value and a flatter slope, separating them from young NT participants. (C) Parameter space spanned by the parameters of the linear regression, the x- (intercept) and y-axis (slope) also separate PPD from NT controls.

### 3.4. Higher Cross-Correlation Between Magnetometer Data in PPD

As a last set of analytics, cross-correlation was performed on each 5 second segment magnetometer data for walking and pointing tasks, and on magnetometer data that were high-pass filtered at 6Hz. When comparing the two tasks – pointing and walking – PPD tend to have more separation from their NT cohorts during the three walking tasks (Figure 14). Their cross-correlation show statistically higher values (χ(1,17) = 7.78, p<0.01) than the pointing tasks(χ(1,17) = 3.78, p=0.05) (see Table B4 for detailed statistical test results). Within these tasks, the most severe PPD exhibits the highest value, while the least severe PPD shows the lowest among the Parkinson’s patient cohort. This result identifies the overall cross-correlations obtained from magnetometer data as the most informative of PPD.

**Figure 14.**
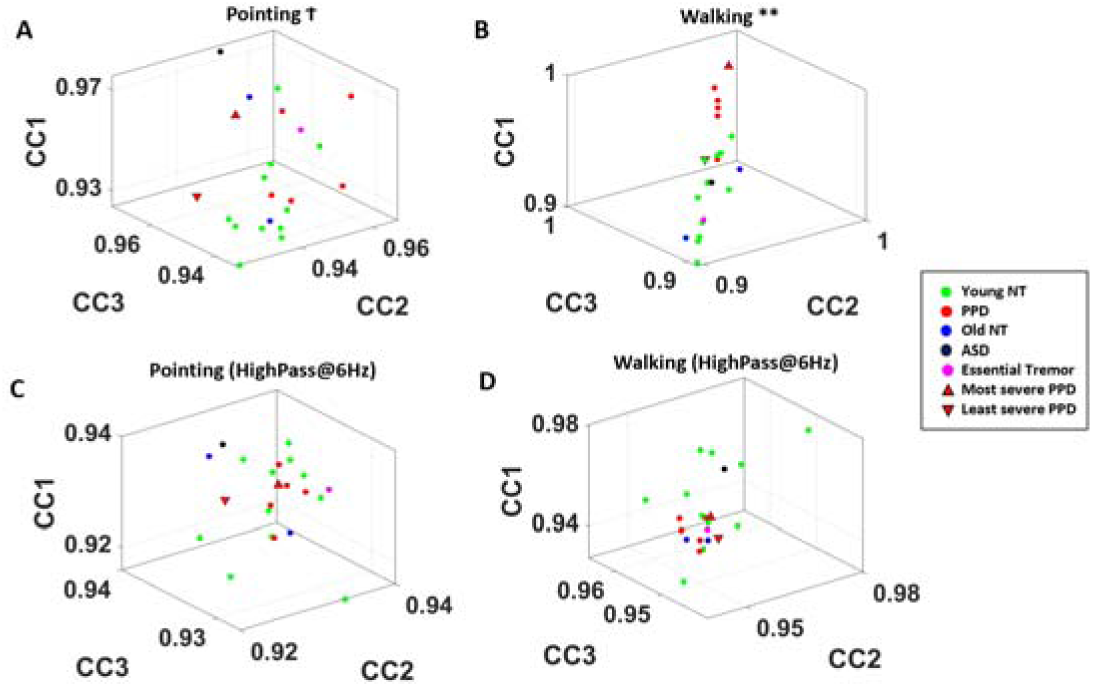
Magnetometer Cross-Correlation in Walking identified as interpretable digital biomarker of PD. (A) Median cross-correlation between all magnetometer sensor pairs for the three pointing tasks (condition P1 in z-axis CC1; P2 in x-axis CC2; P3 in y-axis CC3) are denoted as a single data point for each participant, with colors representing different demographic cohorts. (B) Same as in (A) but for the three walking tasks (condition W1 in z-axis CC1; W2 in x-axis CC2, and W3 in y-axis CC3). Higher cross-correlation in PPD across the different parts of the body reaches statistical significance. Patient with most clinical severity has highest end of the value. (C-D) Same as (A-B) respectively, but for data that were high-passed at 6Hz.

Furthermore, when we examine the clustering pattern of these cross-correlation based on the high-pass filtered data at 6Hz, the pattern no longer exists, and the statistical significance disappears. Moreover, we observe that datapoints of patients with ASD and ET localize within the NT cluster for cross-correlation values of both filtered and un-filtered data.

## 4. Discussion

This work set to investigate multimodal signals from a multiplicity of sensors with different physical units, sampling resolutions and across a heterogeneous cohort of young and elderly NT controls, PPD and other patients with ET and ASD. One motivation of the work was to identify tasks, signals, and parameters that maximally separated PPD from NT controls. Another motivation was to do so while uncovering parameter spaces whereby clinical criteria were conserved, thus lending to higher interpretability of the digital biosensor data. Here the idea is to encourage clinicians to be adopters of the new technology reaching beyond the limits of the naked eye.

Biophysical signals and motor control phenomena are complex and polluted with issues that range from instrumentation noise, the paucity of methods to synchronize activities, the disparity in sampling resolution and interpretable physical units, among others. Further compounding the problem, participants may vary in anatomical sizes possibly confounding kinematic parameters and affecting their variability. Scaling out potential allometric effects and standardizing the time series to reflect significant shifts in the stochastic signatures of the moment-by-moment biophysical fluctuations, led us to the discovery of new features in common, natural tasks amenable to automatically separate PPD from NT controls.

One of the strategies that we used was to study the level of spontaneous entrainment of biorhythmic signals generated during tasks that likely engage primary control from different systems. Among these, we used the tasks of pointing to a visual target and walking. These two tasks were performed while embedded in a context that required disparate levels of intent. These ranged from fully deliberate to spontaneous entrainment of endogenously generated biorhythms with exogeneous rhythmic input. These were accomplished with a metronome that either automatically entrained the biorhythms while playing in the background (no instruction to do so), or intentionally did so by instructing the participant to pace their motions (in pointing) or their breathing (in walking) to the beat of the metronome. These activities evoked statistically significant differentiable responses across multiple signals. However, the walking task did so most pronouncedly in PPD. Furthermore, when examining the various biophysical signals, those from the magnetometer unambiguously separated PPD from NT controls, while conserving the order of severity that traditional clinical tests provided. Along those lines, the patient with the lowest and the one with the highest level of severity revealed themselves on the parameter space as the two extremes. Furthermore, the lowest severity was close to those of the NT controls, while the highest severity was farthest away. This result reveals the magnetometer data, during walking, as highly informative of PD in the digital domain but also as one with high clinical interpretability. In this sense, we render the task and biophysical signal as a potentially interpretable digital biomarker of PD amenable to study at scale using off-the-shelf technology containing this sensor.

A somewhat surprising result in pointing motions is that no significant differences between PPD and NT controls were found when examining the relative distances from the dominant hand to the COM. However, significant differences were revealed in the comparison when examining the left toe fluctuations (contralateral to the moving arm.) Perhaps this alludes to the importance of considering the interplay between spontaneous motions occurring largely beneath awareness, and precision-based motions, requiring deliberate cognitive planning and guidance (*3, 34*). It is possible that the patterns of motor variability here revealed excess uncontrolled manifold motions (*i.e.,* variability in task incidental dimensions linked to CPGs and orthogonal to task relevant dimensions under precision control (*10, 35*)) In this sense, we could think of it as an energy management issue, whereby PPD may exert excess deliberate control of the moving arm and consequentially, have less energy left for the system to monitor or to dampen the noise to signal ratio in lower body end effectors (*e.g.,* the contralateral toe.) This would enhance the contributions from that type of tonic activity from CPG linked variability and broaden the overall statistical departure from controls.

We also examined through the EMD the variability in ISI timing. There we found that the magnetometer, EKG and EEG were the most informative in revealing the largest departure of PPD from NT controls. The PPD showed reduced variability in relation to the NT controls with a far richer range of stochastic shifts depending on the control mode during walking. This was confirmed as well in the Delaunay triangulation analyses related to the EMD scatter derived from the data across all sensors. Lastly, the magnetometer cross-correlation analyses revealed walking as the most informative task differentiating PPD from NT controls. The walking task paired with the experimental assay exploiting different levels of intent and examining the magnetometer data revealed themselves as clinically valid, and optimal in identifying PPD.

### Caveats of the Current Work

A source of limitation in any analyses of brainwaves is the imprecision of the EEG signals. The EEG data is fraught with signals that are not from the cortical processes. The ICA method was applied to alleviate such concern. However, ICA does not allow extracting the same spatial components of EEG signals for all participants (*e.g.,* some participants’ components are mainly from the frontal area, while perhaps other participant’s components may be mainly from the parietal area.) This renders it impossible to get a precise picture and comparison of the brain activities across individuals. The presence of artifacts (in the EEG data) that cannot be clearly identified, further compounds the problem of reproducibility in EEG data. Unfortunately, the current technological state of brain activity measures, does not offer a quick solution to this problem. In our work, the small patient population weakens our characterization of PPD when non-significant patterns emerge. To obtain a clearer picture of such disorder, we would need to collect more data from age-matched control, and other patient populations with compromised motor control. Notwithstanding these issues, the data types, analytical methods, and identification procedure presented here offer new ways to interrogate multimodal data and identify cases where unambiguous separation exists between patients and controls. Furthermore, cases in which such unambiguous separation is paired with good correspondence between clinical criteria and digital data offer the possibility of interpretability and reliable use of the methods in both lab and clinical settings. Such interpretable digital biomarkers could then be adopted by clinicians as a next-generation criteria for features predictive of PD. In this sense, despite limitation, this work could help advance the quest for *interpretable digital biomarkers* of PD and neurotypical aging.

## Author Contributions

Conceptualization, J.R. and E.B.T.; methodology, J.R., R.D., and E.B.T.; formal analysis, J.R. and E.B.T.; investigation, J.R. and E.B.T.; writing—original draft preparation, J.R. and E.B.T.; writing—review and editing, J.R. and E.B.T.; visualization, J.R. and E.B.T.; supervision, E.B.T.; funding acquisition, E.B.T. All authors have read and agreed to the published version of the manuscript.

## Funding

This research was funded by New Jersey Governor’s Council for the Medical Research and Treatments of Autism CAUT17BSP024 and by the Nancy Lurie Marks Family Foundation to E.B.T. Career Development Award.

## Conflicts of Interest

The authors declare no conflict of interest.

## Acknowledgments

The authors declare no financial disclosures. We thank research assistants Christina Wilson, Nishad Nalgundwar

## Data Availability

Data is available upon request to the corresponding author.

## Appendix A. Supplemental Figures

**Figure A1.**
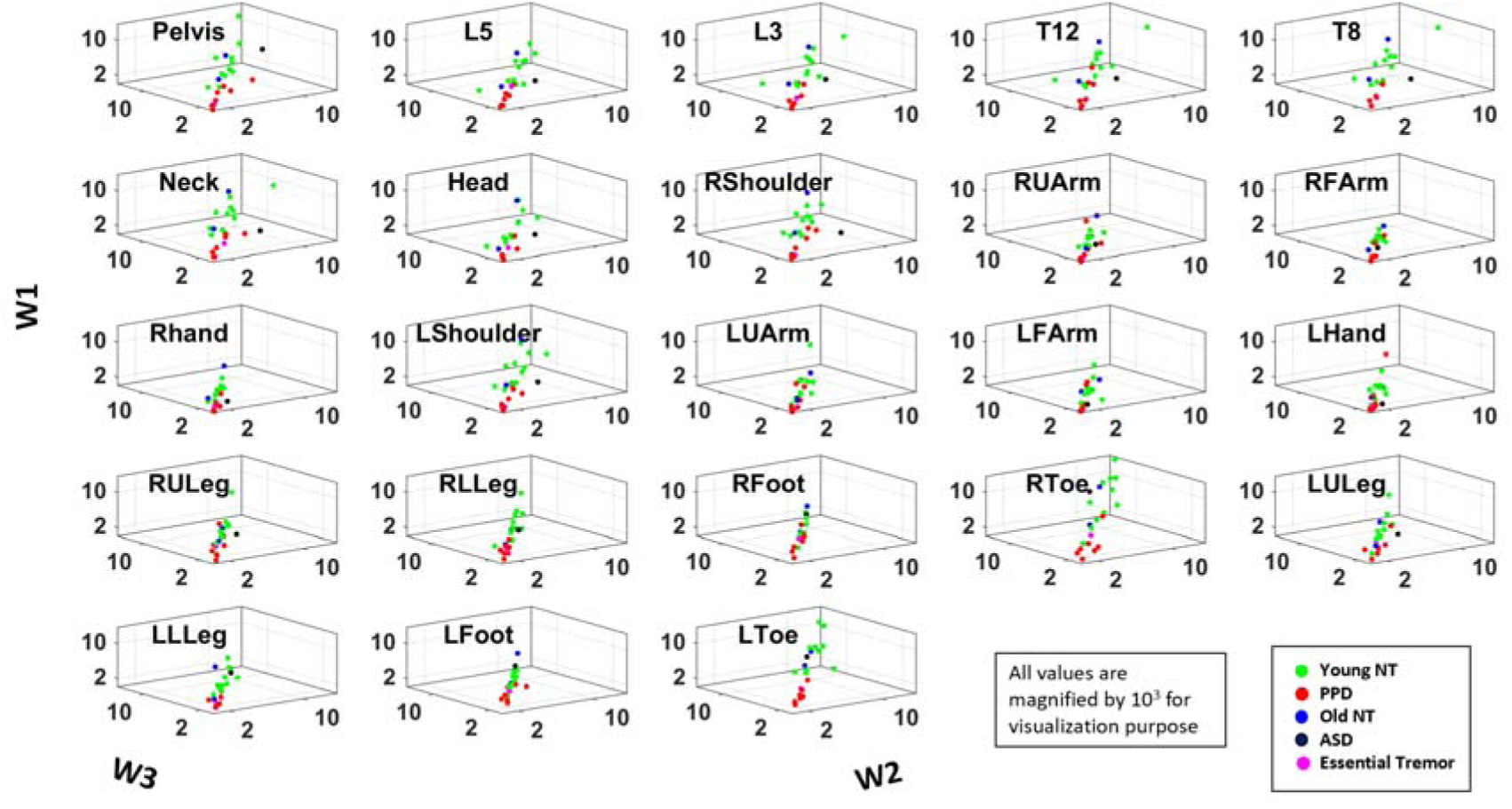
Deviation of the angle between the normal vector and reference vector from π/2 (i.e., abs(angle - π/2), where the normal vector is obtained from planar regression of the 3-dimensional coordinates of distance between body part and COM at time (t,t+1,t+2). The angle deviations are plotted for each body part (subplot) and for each participant with color denoting different demographic cohort during condition W1 (z-axis), W2 (x-axis) and W3 (y-axis).

## Appendix B. Supplemental Tables

**Table B1.** Kruskal Wallis test for angle deviations

| Body Part | Condition | $\chi (1,18)$ | p-value |
| --- | --- | --- | --- |
| Left Toe | W1 | 9.31 | <0.01** |
|  | W2 | 13.00 | <0.01** |
|  | W3 | 13.00 | <0.01** |
| Left Toe | P1 | 3.18 | 0.07 |
|  | P2 | 0.83 | 0.36 |
|  | P3 | 4.41 | 0.04* |
| Dominant Hand | P1 | 0.69 | 0.41 |
|  | P2 | 0.69 | 0.41 |
|  | P3 | 1.71 | 0.19 |

**Table B2.** Kruskal Wallis test for median EMD values

| | $\chi (1,18)$ | p-value |
| --- | --- | --- |
| EEG | 6.63 | 0.01* |
| Mag-EEG | 9.80 | <0.01** |
| Mag-Mag | 11.34 | <0.01** |
| EKG-EEG | 3.39 | 0.07 |
| EKG-Mag | 4.07 | 0.04* |

**Table B3.** Kruskal Wallis Test on EMD (separated by EEG components generated within the brain and those outside the brain)

| | $\chi$ | df | p-value |
| --- | --- | --- | --- |
| IN EEG | 3.08 | (1,14) | 0.08 |
| IN EEG - Mag | 0.54 | (1,14) | 0.46 |
| IN EEG - EKG | 6.88 | (1,16) | <0.01** |
| OUT EEG | 7.05 | (1,18) | <0.01** |
| OUT EEG-Mag | 8.83 | (1,18) | <0.01** |
| OUT EEG-EKG | 3.39 | (1,17) | 0.07 |

**Table B4.** Kruskal Wallis Test on Median Cross Correlation between Participant Cohort

| | $\chi$ | df | p-value |
| --- | --- | --- | --- |
| Pointing | 3.78 | (1,17) | 0.05 |
| Walking | 7.78 | (1,17) | <0.01 ** |
| Pointing HP | 2.07 | (1,17) | 0.15 |
| Walking HP | 0.30 | (1,17) | 0.58 |

## Appendix C. Supplemental Math Explanations

### Appendix C1. Regression onto a plane

Regression onto a plane was done with the 3-dimensional datasets, by computing the distance from each point (x,y,z) to a plane ax + by + cz + d = 0, and finding the coefficients of the plane that minimizes the total distance as such:

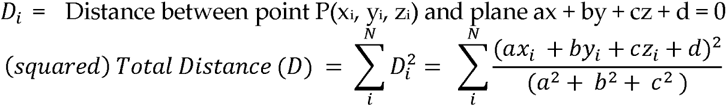

We set the plane equation to reduce one coefficient (d), by taking the derivative with respect to d:

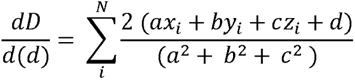

*D* is minimized when we set the numerator (above) to 0, thus providing the following equation:

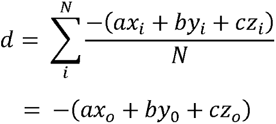

where x_o_, y_o_, z_o_ are means of their respective data points. Using the *d* information above yields the following plane equation:

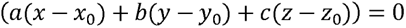

Using the obtained plane equation, we re-formulate the total distance, and minimize this distance as such:

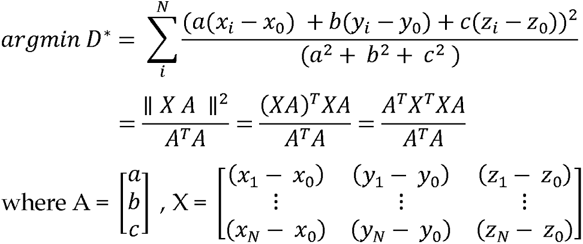

Here, *D\** is represented in a Rayleigh Quotient form and can thus be minimized by the eigenvector corresponding to the smallest eigenvalue of *X^T^X*. Eigenvectors and values were obtained by singular value decomposition (SVD) of *X^T^X*. From the decomposed vectors, the two eigenvectors with larger eigenvalues were set as the axes of the plane, while the eigenvector corresponding to the smallest eigenvalue was set as the normal vector to that plane.

### Appendix C2. Earth Mover’s Distance

The EMD, also known as the Kantarovich-Wasserstein distance (*36*), measures the distance between two discrete probability distributions. Given two discrete distributions P = [(p_1_,w_p1_), … (p_m_,w_pm_)] (*37, 38*), where pi is the cluster representative and w_pi_ is the weight of the cluster; and Q [(p_1_,w_p1_), … (p_n_,w_pn_)], EMD computes how much mass is needed to transform one distribution into another. Defining D [d_ij_] as the ground distance matrix, where d_ij_ is the ground distance between clusters p_i_ and q_j_, and F = [f_ij_] with f_ij_ as the flow between p_i_ and q_j_; EMD is computed by minimizing the overall cost of such:

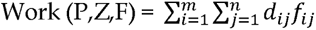

As there are infinite ways to do this, the following constraints are imposed to yield EMD values:

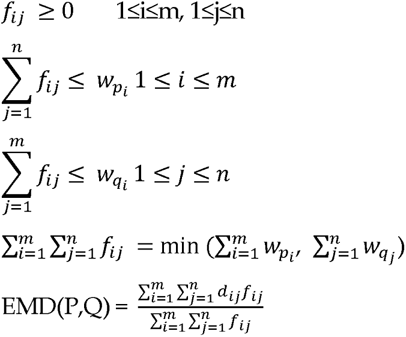

## Notes

### Competing Interest Statement

The authors have declared no competing interest.

